# Anticlustering for Sample Allocation To Minimize Batch Effects

**DOI:** 10.1101/2025.03.03.641320

**Authors:** Martin Papenberg, Cheng Wang, Maïgane Diop, Syed Hassan Bukhari, Boris Oskotsky, Brittany R. Davidson, Kim Chi Vo, Binya Liu, Juan C. Irwin, Alexis Combes, Brice Gaudilliere, Jingjing Li, David K. Stevenson, Gunnar W. Klau, Linda C. Giudice, Marina Sirota, Tomiko T. Oskotsky

## Abstract

High throughput sequencing is a powerful tool for processing large amounts of DNA and RNA samples in batches. Proper experimental design and statistical methods are required to mitigate systematic technical factors due to differences in batches (“batch effects”), as data variation due to these non-biological factors can mask actual biological differences. We propose using anticlustering as an automated method to assign samples to balanced batches. Anticlustering effectively negates differences in (numeric and/or categorical) covariates among batches, and implements user-defined restrictions on the number of batches, the number of samples per batch, and whether to assign certain samples to the same batch (“must-link constraints”). A simulation study shows that anticlustering is better at achieving balance among batches than previous approaches. An application from the UCSF-Stanford Endometriosis Center for Discovery, Innovation, Training and Community Engagement (“ENACT”, https://enactcenter.org/) is presented as a real-life example. In the application, multiple samples provided by an individual had to be processed on the same batch, so that comparisons among different samples of the same patient were not diluted by batch effects. The novel Two Phase Must Link (2PML) anticlustering algorithm realized the must-link restrictions while simultaneously obtaining balance among batches regarding disease stage, menstrual cycle phase, case versus control sample, and clinical site. All methods presented here are accessible via the free and open source R package anticlust (https://cran.r-project.org/package=anticlust). An interactive visualization and web-based batch assignment tool are made available in the Rshiny app “anticlust” (https://anticlust.org/).

## Background/Introduction

With advances in high throughput technologies in recent decades, increasing amounts of data have become available for research, including genomics and epigenomics, bulk, single-cell, and single-nucleus transcriptomics, proteomics, and metabolomics. High throughput sequencing (i.e., “next generation sequencing”), for example, is a powerful method for sequencing large amounts of DNA and RNA that is faster and more cost-efficient than traditional sequencing techniques (e.g., Sanger method), and has contributed to an acceleration of the rate of scientific discovery. Unless the number of samples is sufficiently small to fit in a single batch, samples that undergo high throughput analyses are generally processed in multiple batches. However, systematic technical factors (“batch effects”) can arise when samples are processed in different batches.^1–3^ Proper experimental design and statistical methods are required to mitigate batch effects as data variation due to these non-biological factors can mask actual biological differences.^1–4^ Unless batch effects are appropriately addressed, conclusions from downstream analyses can be misleading or erroneous.^1–3^

Computational packages including sva^5^ and ComBat^6^ can perform batch correction; however, these programs cannot correct all issues with batches. For example, if all samples of patients with a particular condition were assigned to one batch and another batch contained only samples from patients without this condition, then even with batch correction algorithms, it would remain difficult to discern whether differences between the patients’ samples were due to the condition or technical reasons. Hence, it is critical to account for sample variation at the experimental design stage, e.g., avoiding uneven batch sizes, and having balanced classes across batches.^1–3,7^ Resources such as OSAT^8^ and, more recently, a propensity score (PS)-based method^9^ have been proposed to facilitate the allocation of samples to batches so that the impact of batch effects can be minimized. While these approaches have been applied to facilitate real-life batch allocation problems, they come with limitations that reduce their applicability. For example, OSAT is limited to handling categorical variables such as biological sex or race, while numeric variables such as age are not supported (or have to be categorized artificially). The PS-based approach has so far only been implemented to handle up to 4 batches, while real-life applications in high throughput sequencing often have to deal with more batches. Moreover, neither of these approaches implements additional constraints that can be crucial for obtaining valid inferences in experiments. Similar to how teachers want to assign a class of students to equitable groups while keeping some students together in the same group, researchers want to assign samples to balanced batches while keeping some samples (e.g., those from one individual) together in the same batch. There are occasions when researchers want to distribute samples across batches so that the batches have similar representations for features of interest and, at the same time, keep certain samples together on the same batch so that they can be analyzed without concerns of batch effect. For example, if researchers have more than one sample from each patient in their study and they want to compare an individual patient’s samples as well as compare all patients’ samples, then ideally all samples for a particular patient would be on just one batch (not multiple batches) while having balance across the batches for the study. In the context of cluster analysis, these kinds of requirements have been coined must-link constraints^10^, and implementing them can be beneficial in the context of batch assignment.

In this paper, we propose using anticlustering as an automated and improved process to assign samples to balanced batches. Anticlustering is a method that divides a set of elements into disjunct groups, with the aim of maximizing similarity among the different groups.^11–13^ Papenberg and Klau^11^ presented the free and open source R package anticlust that implements a variety of algorithms for anticlustering. Since its introduction, it has been used for diverse tasks in a wide range of research fields.^14–20^ To the best of our knowledge, however, anticlustering has not previously been applied for batch assignment in high throughput sequencing or other high throughput molecular profiling technologies. To close this gap, we provide readers with an introduction to anticlustering, putting a practical focus on batch assignment in high throughput sequencing. Moreover, we introduce a novel method and implementation to include must-link constraints with anticlustering.

To evaluate the usefulness of anticlustering for batch assignment, we conducted an extensive simulation study that compared the anticlust package^11^ to the existing OSAT package and the PS-based method. We demonstrate that anticlust’s ability to assign samples to balanced batches is better than that of these existing methods and anticlust’s must-link constraints did not decrease batch balance considerably. Anticlust batch-assignment and must-link features have already been implemented by the UCSF-Stanford Endometriosis Center for Discovery, Innovation, Training and Community Engagement (“ENACT”, https://enactcenter.org/) for the center’s projects leveraging single-nuclei transcriptomics and other molecular profiling technologies to elucidate mechanisms involved in endometriosis.^21^ Readers who wish to apply anticlustering in their research are provided with additional documentation and code examples in the online supplementary materials (https://osf.io/eu5gd/). All code and data required to reproduce the analyses presented in this paper are openly available from this repository. All anticlustering methods presented here are also openly available and have been implemented in anticlust (https://cran.r-project.org/package=anticlust).

## Results

### Anticlustering algorithm

The standard algorithm anticlust uses to optimize balance among batches is an exchange method that consists of two steps.^11,22^ First, it randomly assigns samples to batches, while adhering to cardinality constraints imposed by the researcher. Our algorithm allows any predefined number of samples per batch, whereas the most common requirement is that each batch must have the same number of samples. The initialization step is followed by an optimization process that systematically exchanges samples between batches. Each exchange is chosen in such a way that it leads to the largest possible improvement in balance (or similarity) among batches. Similarity among batches is quantified through a mathematical objective function that is known from cluster analysis; in cluster analysis, however, samples would be assigned in such a way that the objective function is minimal, while in anticlustering it has to be maximal. We used the popular *diversity* objective, which is based on a mathematical notion of pairwise dissimilarity among individual samples.^12^ In particular, the diversity is computed as the sum of all distances among samples belonging to the same batch (see Figure 1). As the basic distance measure for computing the diversity, we use the Euclidean or squared Euclidean distance based on features, such as age or body mass index (BMI), of the individuals from whom the samples were obtained. Via binary encoding, the diversity can also incorporate categorical variables such as race or disease stage, which are oftentimes considered as parameters when striving for balance among batches.^8,9^ To ensure comparable weight of all features when computing the diversity, it is recommended to standardize the input variables before computing the pairwise distances. Standardization is particularly useful if the variables differ strongly in their ranges.^23^

**Figure 1.**
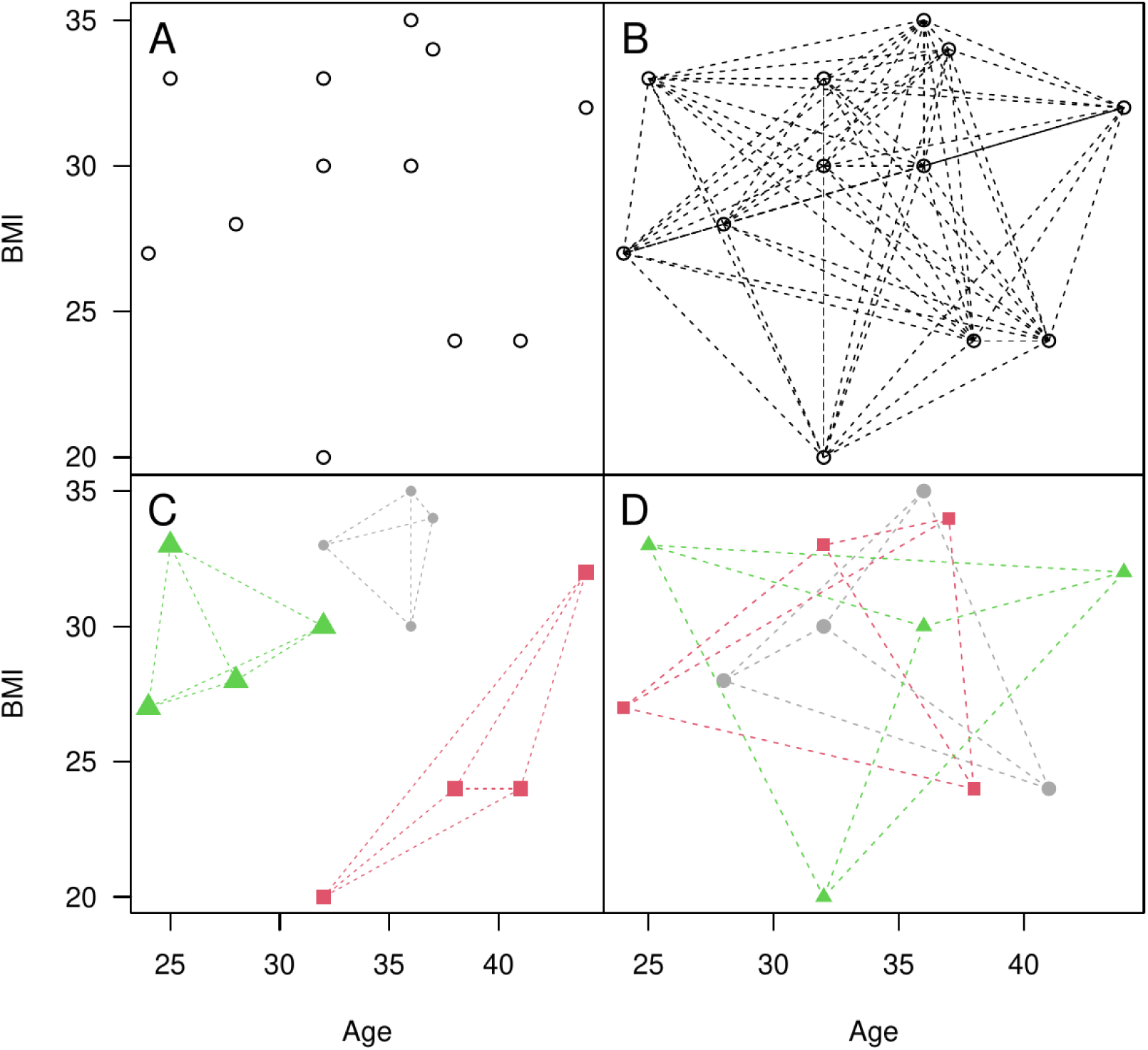
Representation of the conversion from numeric features to Euclidean distance, and (anti)clustering assignments based on minimum and maximum diversity using the Euclidean distance. Panel A illustrates 12 hypothetical values of BMI and age in a scatter plot. Panel B represents the Euclidean distances between features as a straight line in the two-dimensional space. The Euclidean distance is proportional to the length of the connecting lines in panel B. Panel C illustrates a clustering assignment of the 12 data points to *K* = 3 equal-sized groups via *minimum* diversity. Panel D illustrates an anticlustering assignment of the 12 data points to *K* = 3 equal-sized groups via *maximum* diversity. The diversity is computed as the sum of within-cluster distances, which are highlighted in Panel C and Panel D through connecting lines. Maximizing the Euclidean diversity simultaneously leads to similar distribution of the input features among batches (for equal-sized, balanced batches).

To add must-link constraints to anticlustering, we propose a new algorithm called Two Phase Must Link (2PML). Phase 1 is a straightforward adaptation of the standard algorithm for unconstrained anticlustering. To initialize Phase 1, we enforce the must-link constraints through a randomized bin packing algorithm. During the following optimization step, we use a modified representation of the data set in which all samples that must be linked together are treated as a single unit: a *must-link clique*. Using the modified data set, the optimization step applies the same exchange process as the unconstrained algorithm, with the exception that exchanges are restricted to cliques of the same size, thus ensuring that batches remain filled according to the cardinality requirements (e.g., equal-sized batches). Phase 2 drops the restriction of performing exchanges between cliques of the same size and systematically improves over the assignment found in Phase 1. The Methods section describes both phases of 2PML in detail.

### Simulation Study

To thoroughly evaluate the usefulness of anticlustering for batch assignment, we compared it to two alternative approaches in a large-scale simulation study: (1) the OSAT method by Yan et al.^8^, which already has been used frequently for the purpose of batch assignment^24–26^, and (2) propensity score batch assignment (PSBA) by Carry et al.^9^, which has been proposed more recently and has not yet been cited by other researchers. As a secondary contribution of the simulation study, we investigated whether the quality of batch assignment is reduced when must-link constraints are included with anticlustering, as compared to an unconstrained anticlustering assignment. The simulation was implemented using the R programming language (version 4.4.2 ^27^ on an Intel i7-10700 computer (4.800GHz x 8) with 16 GB RAM, running Ubuntu 20.04.6 LTS.

To obtain a fair comparison to the alternative approaches, anticlustering was implemented using the default settings of the anticlust package (Version 0.8.9-1) that—despite numerous updates—have remained unchanged since its original presentation.^11^ That is, we optimized the diversity objective via the exchange method, using the Euclidean distance as the measure of pairwise dissimilarity. Anticlustering subject to must-link constraints used the novel 2PML algorithm, applying one iteration of Phase 1 and one iteration of Phase 2, respectively. For OSAT, we used the implementation that is freely available as an R package from Bioconductor (Version 1.52.0).^28^ We used the default OSAT algorithm *optimal shuffle* with 5,000 repetitions.

We also evaluated the alternative algorithm *optimal block*, but decided not to include it in the final large-scale simulation, because it provided comparable results while running at a significantly slower pace. Carry et al.^9^ provided an R implementation of PSBA that is available from an accompanying online repository (https://github.com/carryp/PS-Batch-Effect). In the original publication, the authors presented a version of PSBA that investigates all possible batch assignments and selects the one that minimizes discrepancy in average propensity scores among batches. However, investigating all possible assignments is only feasible for small data sets and therefore not generally applicable. In the online repository, the authors therefore included an implementation that randomly generates a user-defined number of batch assignments, and selects the best one among them. For comparability with OSAT, we set the number of random assignments to 5,000.

For the simulation, we generated 10,000 data sets. Data sets were processed via (a) OSAT, (b) PSBA, (c) unconstrained anticlustering and (d) anticlustering subject to must-link constraints. Because the OSAT method is only applicable to categorical variables, we generated categorical variables in our simulation. For anticlustering and PSBA, the categorical variables were binary-coded before the methods were applied. OSAT directly used the categorical variables as input because its objective function is computed using category frequencies. For each data set, we randomly determined the number of categorical variables *M* (2-5), the number of classes per variable *P* (2-5), the total sample size *N* (between 50 and 500), and the number of batches *K* (2, 4, or 10 equal-sized batches). PSBA was only applied for *K = 2* and *K = 4* batches because the authors’ implementation only allows the assignment to a maximum of four batches. For constrained anticlustering, must-link constraints were generated by creating a random integer (between 1 and *N*) for each of the *N* samples, which served as an ID variable; that is, if two samples were randomly assigned the same ID, they were required to be assigned to the same batch. This procedure resulted in a distribution of constraints that was similar to the distribution of constraints in our motivating application: 58% of all samples had no must-link partner; 29% had 1 must-link partner; 10% had 2 must-link partners, and 3% had 3 or more must-link partners.

Across the 10,000 simulation runs, the average run time for the competing assignment methods was 0.10s for unconstrained anticlustering, 0.12s for anticlustering with the must-link feature, 3.68s for OSAT, and 31.81s for PSBA, making anticlustering about 29 and 311 times faster than OSAT and PSBA, respectively. For each simulation run for each of the four methods, we computed a χ^2^ test to assess the balance among batches for each of the 2-5 variables, as performed by Yan et al.^8^ In this context, a higher *p*-value indicates that there is more balance among batches, i.e., that the batches are more similar with regard to the covariates. For 84% of all variables that were generated during the simulation, balance among batches was better when using anticlustering as compared to OSAT. Balance was equal in 16% of all comparisons, and only in 0.3% (= 105 of all 34890 pairwise comparisons), OSAT outperformed anticlust. For 78% of all variables that were generated during the simulation, balance was better when using anticlustering as compared to PSBA. Balance was equal in 22% of all comparisons, and only in 0.13% (= 31 of all 23090 pairwise comparisons), PSBA outperformed anticlustering. Figure 2 illustrates average *p*-values in dependence on the number of variables and the number of batches. When increasing the number of variables from 2 to 5, the average *p*-value for OSAT declined from 0.99 to 0.72 whereas the average *p*-value for anticlustering remained greater than 0.99. PSBA also demonstrated a decrease in *p*-value when increasing the number of variables, but less so than OSAT (from 0.99 to 0.91). In the online supplementary materials, we provide Figures illustrating the simulation results while aggregating across the other variables that varied in the simulation (Figure 3: number of samples; Figure 4: number of categories per variable).

**Figure 2.**
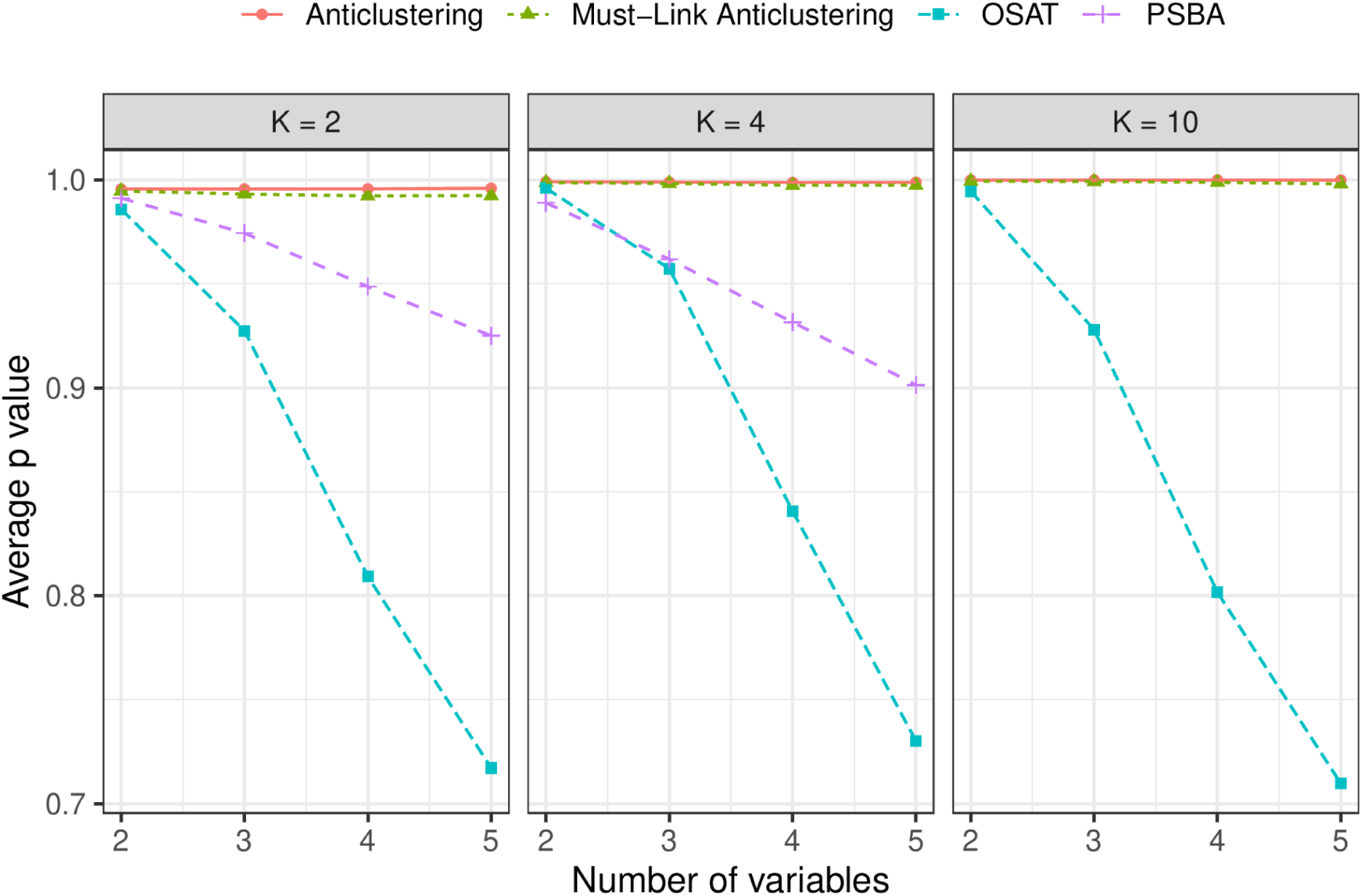
Average *p*-values from χ^2^ tests, based on the number of batches and the number of variables. Higher *p*-values indicate higher balance. Anticlustering maintained a comparable level of balance across all conditions. OSAT’s performance decreased the most with an increasing number of variables.

Remarkably, the constrained anticlustering assignment led to better balance than the OSAT and PSBA methods that did not employ any constraints (see Figure 2). In general, using must-link constraints hardly affected the balance among batches: On average, the anticlustering assignment that was subjected to must-link constraints achieved 99.9% of the objective value of the unconstrained assignment. In an astonishing 75% of all cases, balance was not at all reduced by the must-link constraints. Hence, must-link constraints are not only desirable from a researcher’s point of view, but they also do not contribute to a considerable decrease in batch balance.

### Application

The ENACT Center’s work is concerned with researching endometriosis disease. One of the ENACT projects required assigning 320 tissue samples of women who either have the disease (i.e., case) or not (i.e., controls) to 20 batches (16 samples per batch). To illustrate the application, we prepared a synthetic data set that resembles the actual data set, which is not disclosed for reasons of medical confidentiality. In the synthetic dataset, all women who did not have the disease (*n* = 50) provided exactly one sample, and all women who have the disease (*n* = 89) provided multiple samples: 40 patients provided 2 samples, 25 patients provided 3 samples, 11 patients provided 4 samples, 10 patients provided 5 samples, and 6, 7, or 8 samples were provided by one patient, respectively. In total, the synthetic data set consisted of 320 samples from 139 unique individuals.

For the assignment, we sought balance with regard to four categorical variables: is the disease present (“yes” = case sample; “no” = control sample); stage of disease (“none”, “Stage I or II”, “Stage III or IV”); clinical site (University of California San Francisco (“UCSF”) or other); menstrual cycle phase (proliferative (“PE”) or secretory (“SE”)). Other demographic variables such as age and BMI were also assessed, but not deemed to be critical covariates for the assignment. However, post-hoc checks revealed that no significant discrepancies occurred in these other variables, even though they were not explicitly considered via anticlustering.

In our actual experiment, we only used the results of the must-link restricted assignment. However, for the purpose of illustration, we also applied unconstrained anticlustering and OSAT to show what level of balance can be achieved when no constraints are imposed. Tables 1-3 illustrate the distribution of the categorical variables across batches for unconstrained anticlustering, OSAT, and constrained anticlustering, respectively. The tables were generated using the R package tableone^29^. Table cells contain both information about frequencies and percentages (in parentheses), whether a batch represents only individuals whose samples are unique to that batch (“True” or “False”), and *p-values* from χ^2^ tests for the comparison of the four relevant covariates across the batches. The unconstrained methods successfully balanced all four variables among batches (all *p*-values > .999 for anticlustering; all *p*-values > .99 for OSAT). While all assignments were conducted using the 320 individual samples as input—as opposed to the 139 individual patients—we also verified the satisfactory balance on the level of individuals (see the upper rows in Tables 1-3). Table 3 illustrates the level of balance achieved by constrained anticlustering, which ensured that samples belonging to the same patient were assigned to the same batch. Note that in this application, the must-link constraints put rather severe restrictions on the assignments that were possible; for example, up to 50% of a batch had to be occupied with the samples of a single patient. Still, batches turned out to be highly balanced when including them via the 2PML algorithm (all *p-value*s > .99). Remarkably, our method filled each of the 20 batches with at least one of the 21 individuals with disease Phase I or II (see section Individuals in Table 3).

**Table 1.**
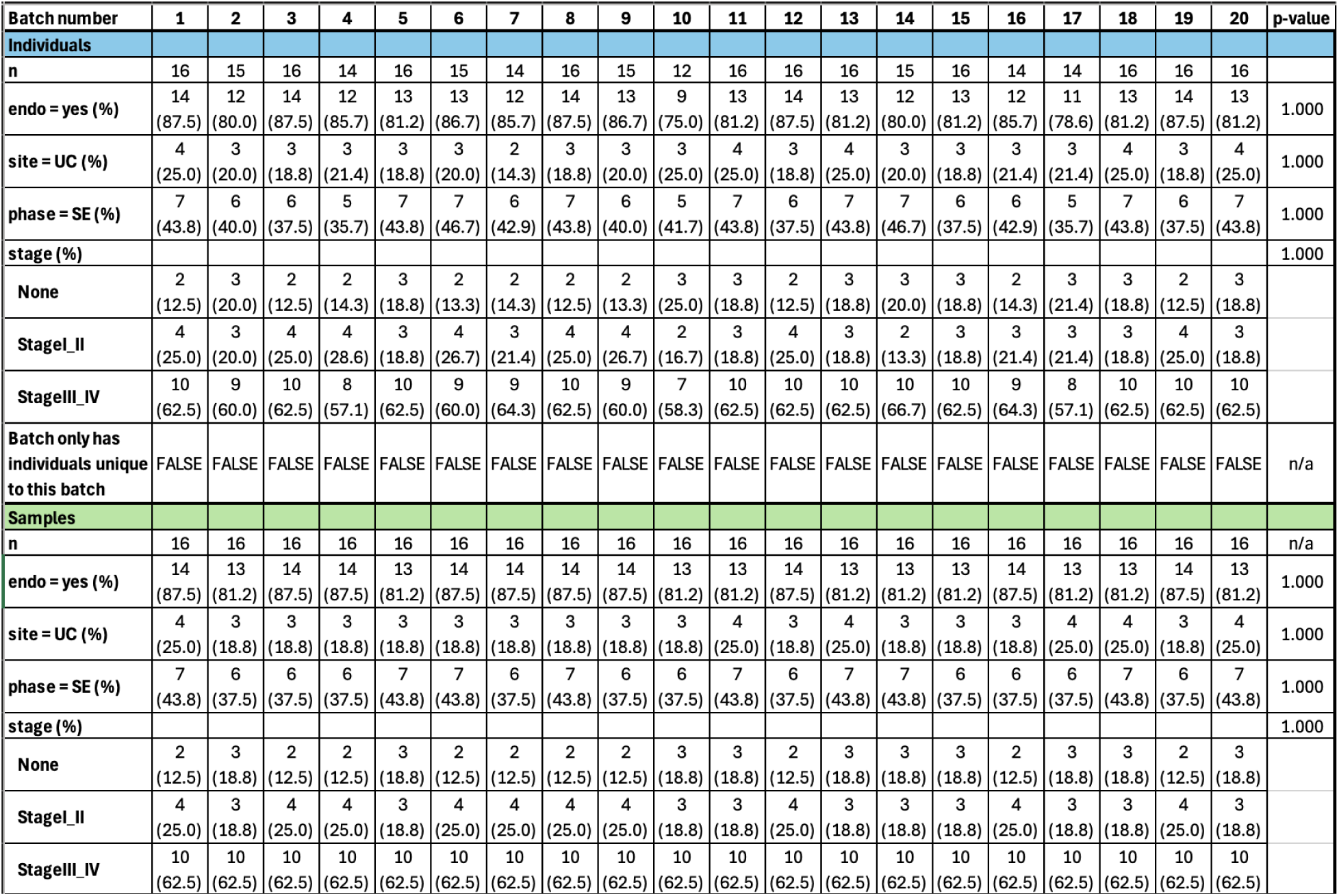
Summary statistics on an individual- and sample level for the batches assigned using the unrestricted anticlustering method, with *p*-values from χ^2^ tests.

**Table 2.**
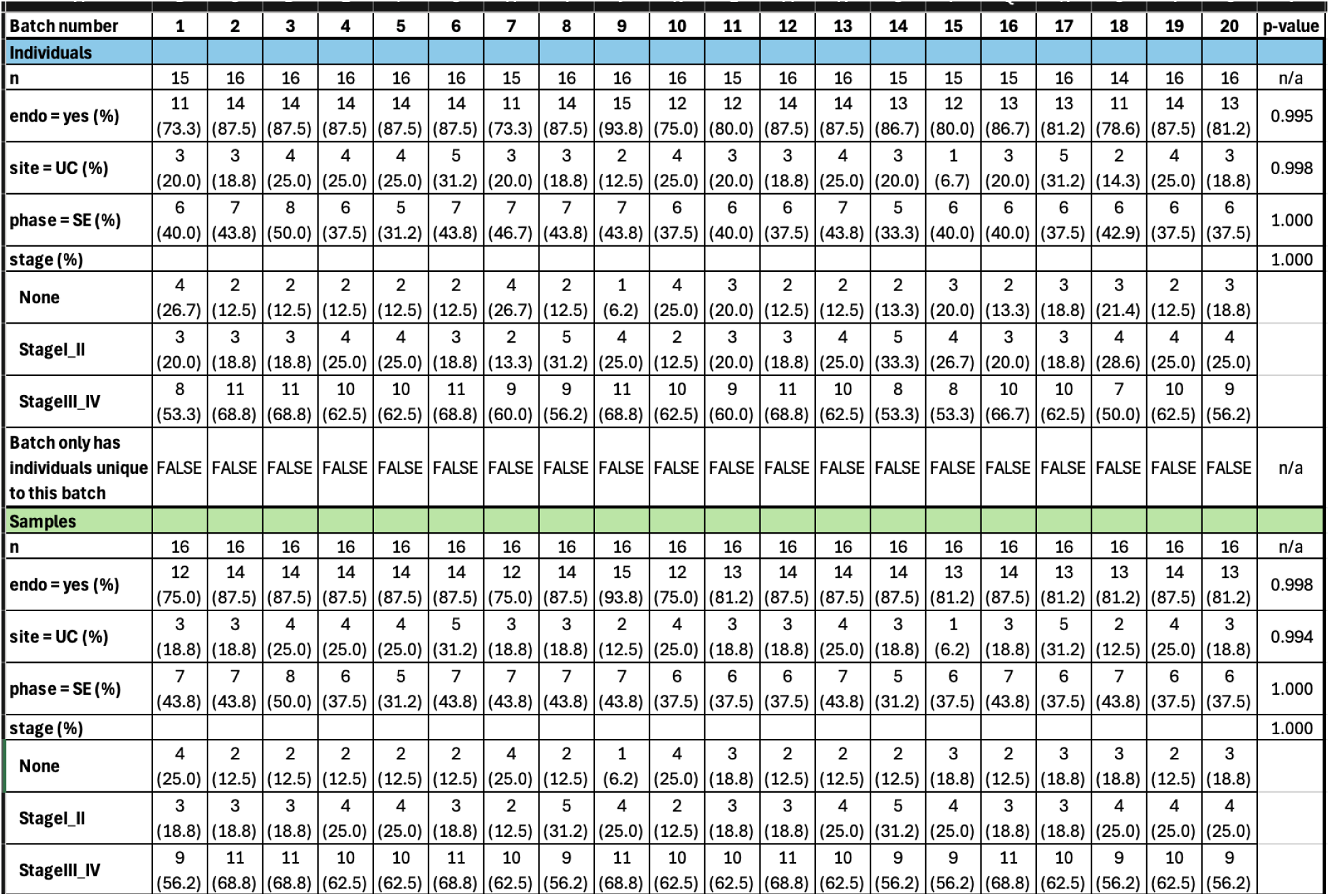
Summary statistics on an individual- and sample level for the batches assigned using the OSAT method, with *p*-values from χ^2^ tests.

**Table 3.**
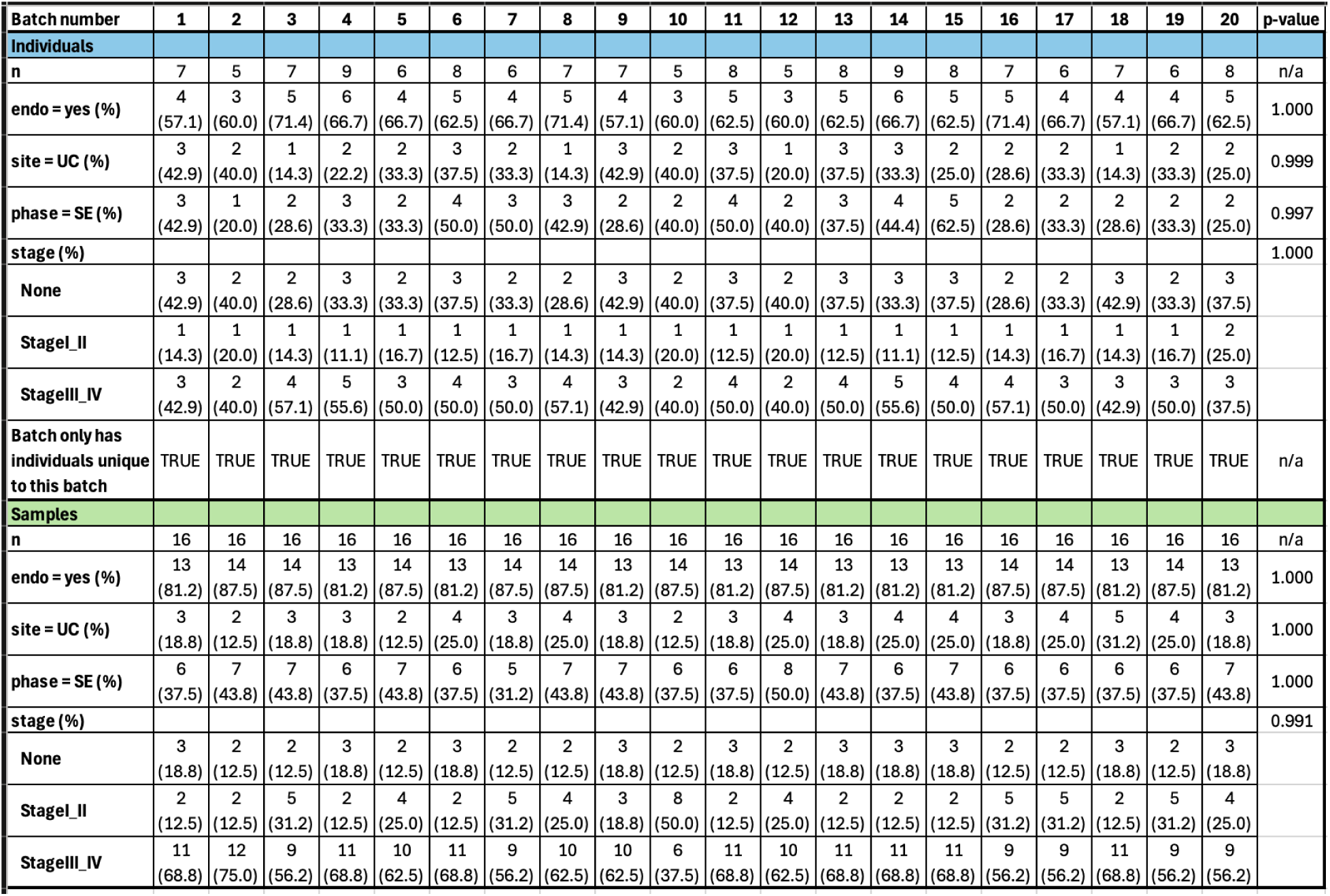
Summary statistics on an individual- and sample level for the batches assigned using the anticlustering method with the must-link feature, with *p*-values from χ^2^ tests.

An interactive visualization of batch assignment for our synthetic dataset with OSAT and anticlust and the balance on a sample level achieved by these methods, as reflected in Tables 1-3, as well as a web-based batch assignment tool, are made available in the Rshiny app “anticlust”, https://anticlust.org/.

## Discussion

High throughput molecular profiling technologies commonly require that samples are processed in batches, while confounding effects among batches should be minimized as much as possible.^8,9^ Though post-hoc corrections during statistical analysis are possible to compensate for imbalances in covariates, exercising control on an a priori level—i.e., creating batches that are as similar as possible on relevant covariates—is crucial as well.^7^ Previous methods for batch assignment include OSAT, which can be used to minimize discrepancy on categorical covariates, and PSBA, which minimizes discrepancy among batches with regard to average propensity scores. We introduced anticlustering as a robust and improved alternative for assigning samples to batches, enabling similarity across categorical and numerical covariates.

While anticlustering has already been used in a variety of settings,^14–20^ it has not yet been put to use in the context of batch assignment, which poses additional challenges as the integration of must-link constraints.

We presented a motivating example to illustrate how anticlustering can be used to effectively assign samples to batches. The example reproduced a real-life application at the UCSF-Stanford Endometriosis Center for Discovery, Innovation, Training and Community Engagement. In our application, we strove for similarity among batches regarding (1) case vs control sample, (2) clinical site, (3) menstrual cycle phase, and (4) disease stage. Anticlustering proved to be an effective tool in order to meet these criteria. We also required that all samples provided by the same person had to be processed in the same batch, to facilitate the comparison of all of an individual’s samples and avoid confounding the analysis with batch effects. For our work, having the ability to assign each individual’s samples intentionally to the same batch would allow greater confidence in findings from our comparison of the individual’s samples as concerns about batch effects would be mitigated. A similar situation is conceivable in the context of assigning students to classes: Groups of friends have to be assigned to the same class, while striving for overall similarity among—and heterogeneity within—classes.

These must-link requirements could not be fulfilled using existing approaches such as OSAT and PSBA, and we therefore developed a novel method (2PML) that for the first time combined the theory of constrained cluster analysis^10^ with the domain of anticlustering.

Our evaluation revealed that anticlustering provides several critical advantages over alternative approaches. First, the simulation study indicates that in the context of achieving balance in batch assignment tasks, anticlustering outperforms OSAT and PSBA quite considerably (Figure 2). Note that these results were not due to a different tradeoff of speed and quality: The improved anticlustering results were obtained much faster than the results of OSAT and PSBA. Moreover, as compared to OSAT and PSBA, anticlustering can be applied more flexibly in a broader range of settings. For example, OSAT’s usage is restricted to using categorical variables as input. Anticlustering can deal with numeric as well as categorical variables, or even using a mixture of these types of variables. Categorical variables can (a) be included as numeric variables via binary coding, as we implemented in the current simulation study and application, or (b) as a “hard constraint” that strictly prioritizes balance according to one categorical variable over the other variables.^11^ PSBA—at least given the current implementation provided by the authors—is restricted to creating a maximum of four batches. Anticlustering has been implemented to handle an arbitrary number of batches, as well as user-defined batch sizes. The anticlust package also provides additional functionality not discussed here, including the possibility to choose among a variety of anticlustering objective functions, different optimization algorithms (including optimal methods), and a specialized algorithm for very large data sets.

The clear superiority of anticlustering over the alternatives OSAT and PSBA, as indicated by our simulation study, might be met with skepticism.^30^ In a strict sense, the results only pertain to the conditions realized in the simulation (e.g., regarding the number of samples or the number and type of variables). Still, we argue that the results reflect a true difference in potency among approaches. In particular, anticlust uses a better optimization algorithm than used by either OSAT or PSBA. The anticlust package by default relies on a variant of the LCW method.^22^ LCW employs a systematic improvement search by exchanging each sample with the most promising candidate that is currently part of a different batch. Anticlust even provides more sophisticated optimization algorithms that have been developed recently, such as the bicriterion iterated local search heuristic by Brusco et al.^12^, and the three-phase search approach by Yang et al.^31^ However, to obtain the most fair comparison in the simulation study, we applied the default method without additional tweaks. Before explaining differences in methodological performance among anticlustering, OSAT, and PSBA in more detail, we would like to clarify an important distinction regarding batch assignment methods. Such methods are generally characterized by two components: (1) an objective function that quantifies balance among batches; (2) an optimization algorithm that assigns samples to batches in such a way that balance is maximized. Among anticlustering, OSAT, and PSBA, both components differ. Hence, observed differences in the simulation study cannot unambiguously be attributed to either component. In particular, decreased performance of OSAT and PSBA does not imply that both components are inferior. Importantly, we believe that the objective functions employed by OSAT and PSBA have merit, but that both methods leave room for improvement regarding their optimization algorithms. First, the PSBA implementation provided by its developers uses a random sampling algorithm. That is, PSBA generates a fixed number of batch assignments at random—possibly while implementing a stratification on one of the input variables—and then selects the one assignment that minimizes discrepancy in propensity scores among samples. While we believe that discrepancy in propensity scores is a useful criterion, mere random sampling is generally considered to be a poor optimization algorithm.^32^ OSAT uses a more advanced optimization algorithm than PSBA, applying pairwise exchanges between samples as does LCW. However, the exchange method used by OSAT is less sophisticated. In particular, it does not conduct a systematic search that selects the best possible exchange for each sample, but instead randomly attempts a (fixed) number of exchanges.

### Limitations

While our evaluation highlights the power of anticlustering for batch assignment, some limitations of the approach should be considered. Anticlustering effectively negates differences in observed covariates among batches, but it is only as potent as the data it receives as input. Whether the covariates anticlustering uses for assignment are actually important to address a research question, is up to the researcher to decide. If important covariates have not been measured, they cannot be balanced. Moreover, anticlustering cannot overcome problems associated with the data themselves. Imprecise measurement and missing data are problems that affect anticlustering just as with any other method of data analysis. The degree to which anticlustering ensures balance also depends on the data. Increasing the number of samples facilitates finding acceptable balance; using additional covariates or smaller group sizes reduces balance among batches.^23^ Given that these parameters usually cannot be chosen freely, researchers should always carefully inspect the observed balance after applying anticlustering. Another limitation of the anticlustering approach is that it can only be used if all samples are available before conducting batch allocation; anticlustering requires that all covariates have been measured at the start. It cannot be used to sequentially fill batches as additional samples become available.^33^

## Conclusion

In this paper, we introduced anticlustering as an effective method to assign samples to batches to mitigate batch effects and ensure the robustness of analyses of data from next-generation sequencing and other high throughput molecular profiling technologies. Additionally, we implemented a novel feature for anticlustering, which allows researchers to specify pairs or groups of samples that must be assigned to the same batch. Anticlustering including its must-link feature is freely available via the open source R package anticlust (https://CRAN.R-project.org/package=anticlust) and can be interactively visualized and utilized through the R Shiny app, https://anticlust.org. Via the accompanying package website (https://github.com/m-Py/anticlust), additional documentation and community support are available.

## Methods

Anticlustering is an optimization method that is characterized by (a) an objective function that quantifies the balance among batches, and (b) an algorithm that conducts the batch assignment in such a way that balance among batches is maximized. Anticlustering owes its name to the fact that the objective functions it uses are the reversal of criteria used in classical cluster analysis. For example, Späth^13^ already recognized that by maximizing instead of minimizing the k-means criterion (the “variance”), he was able to create groups that are similar to each other, and presented it as an improvement over the more intuitive random assignment.^34^ Brusco et al.^12^ recognized that other objective functions known from cluster analysis can also be implemented in the context of anticlustering.

In our application, we optimized the *diversity* objective to maximize similarity among batches. While diversity is technically a measure of within-batch heterogeneity, its maximization simultaneously leads to minimal difference between the distribution of the input variables among batches (at least when batches are equal-sized).^23,35^ Papenberg and Klau^11^ referred to the maximization of the diversity as “anticluster editing” because the minimization of the diversity is also well-known from the area of cluster analysis—under the term “cluster editing”.^36,37^ The diversity is computed on the basis of a measure of pairwise dissimilarity among samples. In particular, it is defined as the overall sum of all dissimilarities among samples that are assigned to the same batch.^12^ Hence, the diversity is not directly computed on the basis of sample features, but instead it relies on a distance measure that is computed on the basis of these features, for each pair of samples. In the context of anticlustering, the Euclidean distance is the most common measure that translates features to pairwise dissimilarities.^11,38^ The Euclidean distance is defined as

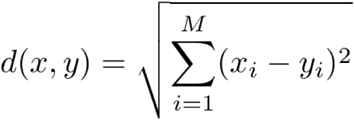

where *M* is the number of numeric features describing two samples *x* = (*x*_1_, …, *x_M_*) and *y* = (*y*_1_, …, *y_M_*). When samples are described by two features, the Euclidean distance corresponds to the geometric, “straight line” distance between two points in a two-dimensional space; more similar data points are closer to each other (see Figure 1). For categorical variables, we use one-hot binary encoding^39^ before including them in the computation; in our example application, we used the squared Euclidean distance on binary encoded features as input for anticlustering.

An anticlustering algorithm assigns samples to batches in such a way that the objective function—here, the diversity—is maximized. Anticlustering usually employs heuristic optimization algorithms.^31^ While heuristics generally provide satisfying results in the context of anticlustering^11^, they do not guarantee finding the globally best assignment among all possibilities. In principle, enumerating all possible assignments is a valid strategy to obtain an optimal assignment. However, this approach quickly becomes impossible due to an exponential growth of the way in which assignments can be conducted.^11^ Moreover, because anticlustering problems (with the exception of some special cases) are also NP-hard^35^, there is likely no algorithm that identifies the globally best assignment without considering all possibilities (at least in the worst case). In practice, heuristics are therefore indispensable.

In the anticlust package, a heuristic exchange algorithm is by default used to maximize the diversity.^11,13,22^ It consists of two steps: an initialization step and an optimization step. As initialization, it randomly assigns samples to batches of equal size or user-defined sizes. After initialization, the algorithm selects a sample and checks how the diversity would change if the sample were swapped with each sample that is currently assigned to a different batch. After simulating each exchange, it realizes the one exchange that increases the diversity the most. It does not conduct an exchange if no improvement in diversity is possible. This procedure is repeated for each sample and it terminates after the last sample is processed. The procedure might also restart at the first element and reiterate through all samples until no pairwise exchange can lead to any further improvement, i.e., until a local maximum is found. In anticlust, we also implemented this local maximum search, which corresponds to the algorithm LCW.^22^ For better results, it is also possible to restart the search algorithm multiple times using different (random) initializations.^13^

### Two Phase Must Link Algorithm

In our application, samples belonging to the same patient were required to be assigned to the same batch. We refer to a set of samples that must be linked together within the same batch as a must-link *clique*. We use the term *singleton* to refer to samples that are not part of a clique.

Basically, anticlustering with the must-link feature uses a modified representation of the original data set in which singletons and cliques are used as input elements—as opposed to unconstrained anticlustering, where each individual sample constitutes an input element. To implement the must-link requirements while striving for similarity among batches, we developed the Two Phase Must Link (2PML) algorithm for anticlustering.

#### Phase 1

The first phase of 2PML is an adaptation of the local maximum search algorithm LCW, which however ensures that must-link constraints are respected. Some adjustments of the algorithm are required to ensure (a) we obtain a valid partitioning regarding the cardinality constraints (e.g., equal-sized batches) and (b) the diversity is computed correctly during optimization. We therefore had to adjust both the initialization phase as well as the optimization phase of the algorithm.

During initialization, we first assign all cliques to batches. Each clique must be assigned completely to one of the batches and samples within a clique must not be split apart. At the same time, the maximum capacity of each batch must not be exceeded. Using this conceptualization, the initialization step corresponds to a bin packing problem, which is one of the classical NP-complete problems in computer science.^40^ That is, we assign a weight to each clique, corresponding to the number of samples it contains. When filling batches, the sum of the weights of the cliques in each batch must not exceed its capacity. Many exact and heuristic bin packing algorithms have been developed. As the default method, we use a *randomized first fit* heuristic to fill batches: For each clique, we iterate through all batches in random order and assign it to the first batch where it fits. The process is expected to evenly distribute the must-link cliques among batches. This random component is particularly useful if we apply multiple restarts of the optimization algorithm. After assigning cliques to batches, the singleton samples can be assigned arbitrarily to fill the remaining space. Note that our randomized first fit algorithm is a heuristic that may not find an assignment of must-link groups to batches even if one is theoretically available. If the heuristic indicates that the batches cannot hold the must-link cliques, we therefore use an exact algorithm based on integer linear programming (ILP) as a fallback option, which allows us to verify if the constraints really cannot be satisfied. To this end, we implemented an adaptation of the standard bin packing ILP model by Martello and Toth^41^:

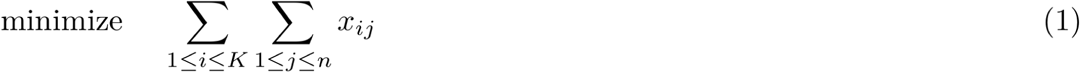

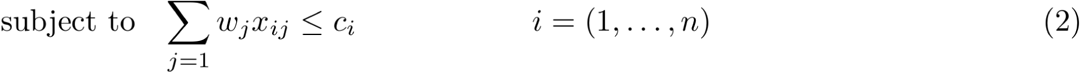

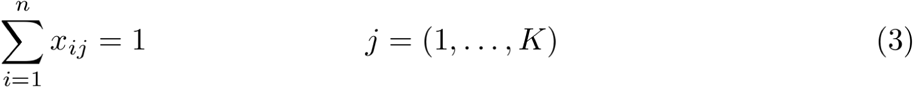

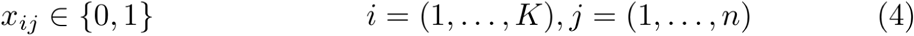

The number of cliques is given by *n*. The model has binary decision variables *x_ij_* to encode whether clique *j* (*j = 1, …, n*) is assigned to batch *i* (*i = 1, …, K*). The model uses *K* values *c_i_* to represent the capacity of each batch. The model has *n* values *c_j_*to encode the weight of each clique, i.e., the number of samples it represents. Constraint (2) ensures that the weight of each batch is not exceeded; constraint (3) ensures that each clique is assigned to exactly one batch. Note that during the initialization step, we only need to test if the constraints (2) and (3) can be satisfied; in the context of must-link anticlustering, any feasible assignment is equally valid. For this reason, the objective function (1) is chosen to be constant for each assignment that satisfies the constraints. It does not actually contribute to solving the problem, and the model therefore only tests if the must-link constraints can be satisfied. In anticlust, we call one of the programs Symphony^42,43^, GLPK^44^, or lpSolve^45^ to yield an optimal assignment for the bin packing problem as initialization for the must-linked samples.

For the optimization phase of the exchange algorithm, we generate a modified representation of the original distance matrix that preserves all relevant information. To this end, we compute new pairwise distances between (a) each clique and the other cliques and between (b) each clique and the singleton samples. The new distances are given as the sum of all pairwise distances between (a) the samples in two different cliques, or (b) between all samples of a clique and a singleton. In the context of maximizing the diversity, this transformation sufficiently preserves the relevant information in the original distance matrix.^37^ Using the initial assignment and the transformed distance matrix, we apply the same exchange algorithm as we described for the unrestricted anticlustering problem. However, during the exchange process, we only permit exchanges between cliques of the same cardinality (e.g., patients providing the same number of samples) to ensure that the cardinality constraints are respected throughout. As opposed to unconstrained anticlustering, this restriction limits the number of exchanges that are evaluated. Hence, we developed Phase 2 to overcome the limitations of Phase 1.

#### Phase 2

The second phase of 2PML is an improvement phase that drops the restriction of performing exchanges among must-link cliques of the same cardinality. It is initialized using the assignment obtained in Phase 1. It iterates through all must-link cliques and for each clique, selects a combination of samples for exchange. The combination of samples is chosen in such a way that (a) must-link constraints are preserved, (b) all samples are currently part of the same batch, and (c) the total number of samples is equal to the size of the current clique. Among all combinations of samples that fit these requirements, one combination is randomly chosen. If no feasible combination is found, the next clique is evaluated. If a combination is found that fits the requirements (a)-(c), the effect of exchanging it with the current clique is evaluated. Exchanges that do not lead to an improvement in diversity are discarded; exchanges that lead to an improvement in diversity are retained. For example, for a clique of size 4, we might select four singletons for exchange; a combination of two cliques of size 2; a clique of size 3 and a singleton; or two singletons and a clique of size 2. Note that generating all possible combinations for exchange corresponds to the subset sum problem, which is another classical NP-hard problem in computer science.^40^ Due to its computational complexity, we cannot exhaustively generate all exchange combinations for large clique sizes. We exhaustively generate all combinations for cliques of up to 10 samples; for larger cliques, we use a heuristic to generate a reduced set of feasible exchange combinations: We generate a set of equal-sized cliques, possibly filled with singletons to ensure the number of samples is sufficient. For example, if this rule were applied to cliques of size 7 (which it is not, because we generate all possible combinations for cliques of size 7), we would obtain the combinations (1, 1, 1, 1, 1, 1, 1), (2, 2, 2, 1) and (3, 3, 1) for exchange.

After iterating through all cliques, the improved batch assignment is once again used as input for the optimization step of Phase 1. That is because—maybe counterintuitively—while the new assignment is likely improved over the initial assignment that was obtained in Phase 1, it may no longer be locally optimal. It can therefore be restored to local optimality for additional improvement. The idea of restoring local optimality of a perturbed assignment is inspired by the iterated local search by Brusco et al.^12^

#### Multiple Restarts

Both phases of 2PML can be called repeatedly as part of a multiple restart algorithm. In anticlust, when users request multiple restarts, half of the restarts will perform Phase 1 and the other half will perform Phase 2. The best assignment across all restarts in Phase 1 is used as input for the first iteration of Phase 2. The second iteration of Phase 2 will again perform an improvement phase, this time using the output of the first iteration of Phase 2 as input; the third iteration then uses the output of the second iteration, and so forth. The process is repeated until all user-defined restarts are conducted.

### Optimal anticlustering using must-link constraints

Due to the computational complexity of anticlustering, batch assignment problems are usually tackled using heuristic algorithms. Still, for some problem constellations—in particular when *N* is not large—it is possible to employ exact algorithms that find the globally best batch assignment.

Papenberg and Klau^11^ presented an ILP model to find globally optimal batch assignments for the diversity objective. It can be used to solve problem instances of up to about *N = 30* in acceptable running time; Schulz^46^ used a time limit of 1800 seconds and showed that up to 30 samples can be assigned optimally in this time. In the current paper, we extend the model by Papenberg and Klau^11^ to allow it to include must-link constraints. The extension is actually quite straightforward: To induce must-link constraints in the context of an exact algorithm, it is sufficient to adjust the distance matrix used as input. Whenever the pairwise distance between two samples is set to ∞—and the set of must-link constraints can be satisfied—any globally optimal assignment will place these samples in the same batch, because the objective value associated with such an assignment is necessarily better than that of an assignment that places them in different batches. This “trick” of adjusting the data input to induce must-link constraints has long been used in the context of exact algorithms for cluster editing.^37^ Including must-link constraints in anticlustering increases the number of samples that can be processed, because it reduces the effective number of samples. In our online supplementary materials, we show that up to about 40 samples can be processed using a time limit of 1800 seconds on a desktop computer; beyond that, the running time is expected to increase exponentially with an increasing number of samples.

## Acknowledgements

This study was funded, in part, by the National Institutes of Health, the Eunice Kennedy Shriver National Institute for Child Health and Human Development, P01 HD106414. The funder played no role in study design, data collection, analysis, and interpretation of data, or the writing of this manuscript. The authors sincerely thank Martin Breuer, Jessica Gelfand, Hannah Hengelbrock, Edna Rodas, and Vivian Siu for their valuable contributions to this work.

